# Slow fluctuations in ongoing brain activity decrease in amplitude with ageing yet their impact on task-related evoked responses is dissociable from behaviour

**DOI:** 10.1101/2021.11.18.467545

**Authors:** Maria J. Ribeiro, Miguel Castelo-Branco

**Affiliations:** Coimbra Institute for Biomedical Imaging and Translational Research (CIBIT), Institute for Nuclear Sciences Applied to Health (ICNAS), University of Coimbra, Portugal; Faculty of Medicine, University of Coimbra, Portugal

**Keywords:** EEG, pupillometry, ageing, brain variability, reaction time variability

## Abstract

In humans, ageing is characterized by decreased brain signal variability and increased behavioural variability. To understand how reduced brain variability segregates with increased behavioural variability, we investigated the association between reaction time variability, evoked brain responses and ongoing brain signal dynamics, in young (N = 36) and older adults (N = 39). We studied the electroencephalogram (EEG) and pupil size fluctuations to characterize the cortical and arousal responses elicited by a cued go/no-go task. Evoked responses were strongly modulated by slow (< 2 Hz) fluctuations of the ongoing signals, which presented reduced power in the older participants. Although variability of the evoked responses was lower in the older participants, once we adjusted for the effect of the ongoing signal fluctuations, evoked responses were equally variable in both groups. Moreover, the modulation of the evoked responses caused by the ongoing signal fluctuations had no impact on reaction time, thereby explaining why although ongoing brain signal variability is decreased in older individuals, behavioural variability is not. Finally, we showed that adjusting for the effect of the ongoing signal was critical to unmask the link between neural responses and behaviour.

## Introduction

Brain signal variability holds valuable information about brain dynamics that should not be ignored when studying the link between neuronal activity and cognition. Brain signal variability can take many forms. Moment-to-moment variability of the ongoing signal reflects how much brain activity changes from one moment to the next, i.e., the brain activity range. Trial-by-trial variability in task-related evoked responses reflects differences in the brain responses to repeated task conditions, while brain signal entropy quantifies the irregularity of the time series.

Within-subject brain signal variability has been shown to change with ageing and is a robust marker of age (Grady & Garrett, 2014). The effect of ageing on brain signal variability has been studied using several measures, including temporal standard deviation (SD) and mean square successive difference (MSSD) of the blood-oxygen-level-dependent (BOLD) signal acquired with functional magnetic resonance imaging (fMRI) (Grady & Garrett, 2014; Samanez-Larkin et al., 2010), temporal SD of the spectral power within frequency bands measured with electroencephalography (EEG) (Kumral et al., 2020), entropy in EEG and magnetoencephalography (MEG) signals (Kosciessa et al., 2020; McIntosh et al., 2014; Waschke et al., 2017), the slope of the EEG power spectral density (PSD) (Voytek et al., 2015), and the consistency of task-related evoked responses (Sander et al., 2012; Tran et al., 2020). The temporal standard deviation of the BOLD signal and of the amplitude envelop of EEG frequency bands measured in the ongoing signal are markedly decreased in older adults (Garrett et al., 2015; Grady & Garrett, 2018; Kumral et al., 2020). Yet, the localization of these changes does not coincide suggesting that BOLD and EEG variability capture different aspects of brain function. In fact, age differences in BOLD SD might be explained by changes in cardiovascular and cerebrovascular factors that affect the link between neuronal activity and changes in blood flow (Tsvetanov et al., 2020). Moreover, BOLD MSSD, a measure of how much the BOLD signal changes across successive observations, is more often increased rather than decreased in older adults (Boylan et al., 2021; Nomi et al., 2017; Samanez-Larkin et al., 2010), highlighting the fact that age-related changes in signal dynamics can result in more or less “signal variability” depending on the used metric. In fact, the relationship between the non-invasive measures of brain signal variability used in human neuroscience and neuronal noise is not trivial. Large signal fluctuations will increase signal standard deviation but might originate in a more stable and predictable signal associated with reduced noise. Higher signal variability is, therefore, not equal to a noisier signal. Consistent with this idea, EEG signal entropy increases with ageing and is associated with the EEG spectra slope (Waschke et al., 2017). Older adults present EEG signals with flatter spectra, i.e., more similar to white noise, potentially reflecting increased neuronal noise (Voytek et al., 2015). Higher brain noise is also consistent with the fact that older people present increased behavioural variability measured as less consistent reaction time across trials (Grady & Garrett, 2018; MacDonald et al., 2012).

However, the link between brain signal variability, neuronal noise and behavioural variability remains to be clarified.

The ongoing brain signal (*i.e.* spontaneous brain activity) measured with fMRI or EEG during “task-free”, “resting” periods reflects activity in large-scale networks each identified as a set of brain regions that show activity co-fluctuations, the resting-state networks (Beckmann et al., 2005; Liu et al., 2017). Notably, these networks are functionally active also during task performance (Abreu et al., 2020; Smith et al., 2009). The ongoing activation and de-activation of these networks presents a backdrop of activity on top of which task-related activity occurs (Fox et al., 2006). Not surprisingly, ongoing activity modulates task-related evoked responses that in turn affect task performance (Becker et al., 2011; Fox et al., 2007; Mayhew et al., 2013; Ribeiro et al., 2016). For example, fluctuations in pre-stimulus alpha power affect the amplitude of brain sensory responses, perception and reaction time (Lou et al., 2014; Thut, 2006; van Dijk et al., 2008). Older people display low-frequency (<12 Hz) brain signal fluctuations with reduced amplitude (Kumral et al., 2020), however, previous studies of the effect of ageing on the variability of task evoked responses have yielded contradictory results, revealing increased or decreased trial-by-trial consistency depending on stimulus or task characteristics (Sander et al., 2012; Tran et al., 2016, 2020; Wiegand & Sander, 2019). Finally, to our knowledge, no previous study has investigated the effect of ageing on the link between the dynamics of the ongoing brain activity, trial-by-trial variability of evoked responses and behavioural variability.

In this study, we aimed at clarifying the effect of ageing on brain signal dynamics and its impact on behaviour by investigating the relationship between variability of ongoing brain signals, evoked responses, and behaviour. We analysed EEG and pupil data from a previously published study acquired while a group of young and a group of older adults were engaged in a cued auditory go/no-go task (Ribeiro & Castelo-Branco, 2019a, 2019b, 2021). The EEG allows the non-invasive measurement of cortical electrical activity with high temporal resolution, while the pupillogram, acquired under conditions of constant luminance, is associated with fluctuations in arousal and reflects activity in the ascending brainstem neuromodulatory systems (Gee et al., 2017; Joshi et al., 2016; Murphy et al., 2014; Reimer et al., 2016). Although, activity modulation in subcortical arousal nuclei (reflected in changes in pupil size) correlates with the EEG signal (*e.g.* Hong et al., 2014; Podvalny et al., 2021), these two signals capture distinct aspects of brain function, both important and complementary for our understanding of the ageing brain. In our previous study, we showed that, in both signals, the cue stimulus evoked a preparatory response. In the EEG, the cue evoked a frontocentral negative potential [the contingent negative variation (CNV)] that increased in magnitude throughout the preparatory period. The CNV has been previously shown to correlate with reaction time on a trial-by-trial basis (Boehm et al., 2014), suggesting that trial-by-trial amplitude fluctuations in the CNV are associated with behavioural variability. In the pupillogram, the cue evoked a pupil dilation response of equal magnitude in both groups of participants (Ribeiro & Castelo-Branco, 2019a). In the current study, we investigated how trial-by-trial variability in the amplitude of these preparatory responses changed across age groups, how it related to behaviour variability and how it was affected by ongoing brain signals. We found that, while behavioural variability was increased in older adults, this group of participants presented reduced variability of the evoked responses both in the EEG and in the pupillogram. We found that the decreased variability in the evoked responses emerged due to a reduction in the amplitude of the slow fluctuations observed in the ongoing signals and did not reflect reduced variability of the evoked responses per se. Once differences in the dynamics of the ongoing signals were accounted for, we observed no group differences in the variability of the evoked responses. Moreover, although the evoked responses were markedly affected by fluctuations in the ongoing signals, the amplitude modulations caused by the ongoing signal did not contribute towards behavioural variability, thereby explaining why although ongoing brain signal variability is decreased in older individuals, behavioural variability is not.

## Results

We analysed EEG and pupil data acquired while a group of young (N = 36; age = 23 ± 3 years) and a group of older (N = 39; age = 60 ± 5 years) adults were engaged in a cued-auditory go/no-go task (Figure 1A).

**Figure 1.**
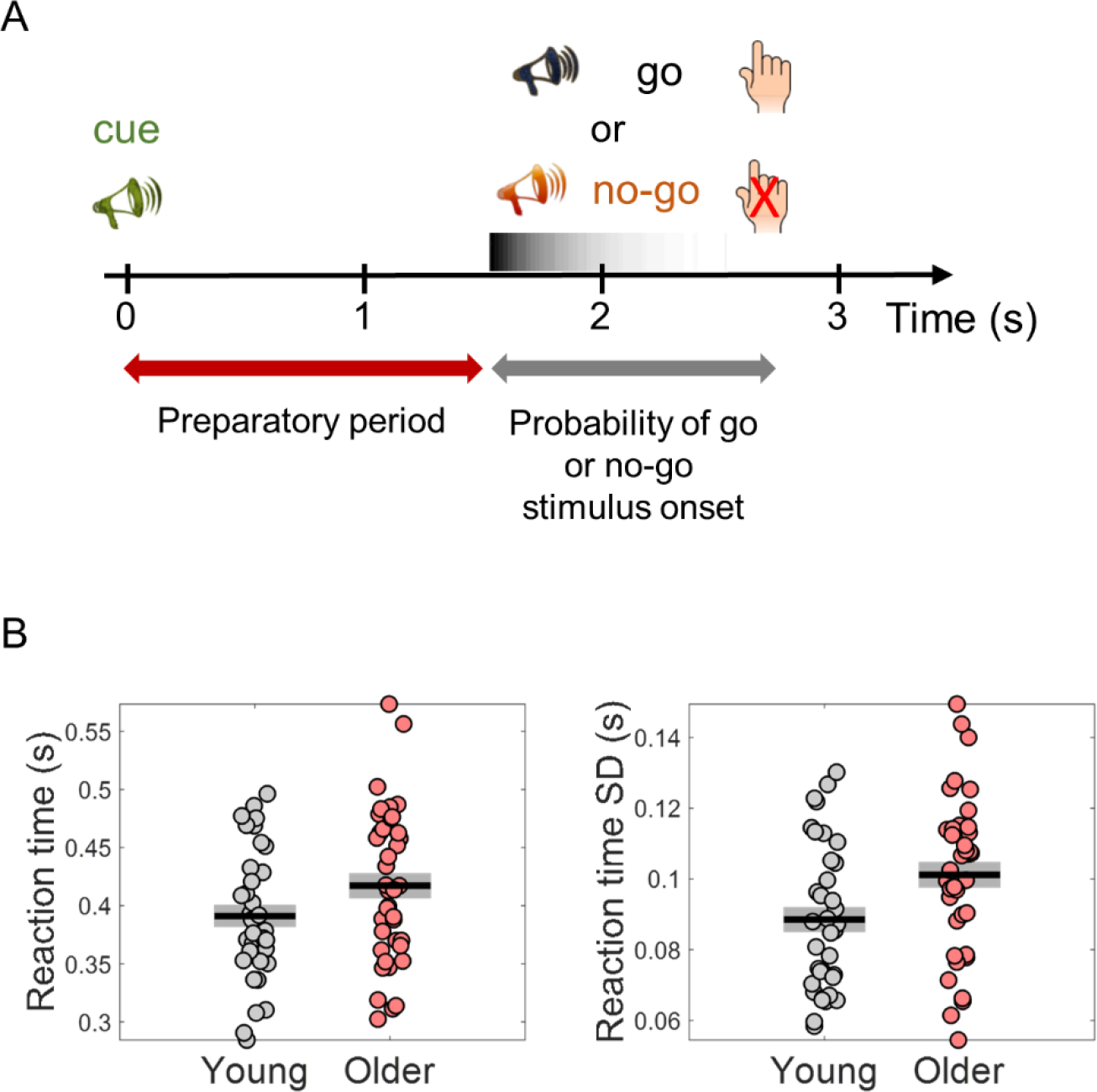
Behavioural task design and reaction time results. (A) Participants performed a cued auditory go/no-go task. The warning cue presented at the beginning of the trial was followed by a go or a no-go stimulus. Participants were instructed to respond with their right index finger as fast as possible upon detection of the go stimulus and to withhold the response in no-go trials. Auditory stimuli were pure tones of different frequencies (cue – 1500 Hz; go – 1700 Hz; no-go – 1300 Hz). (B) Median reaction time (left) and reaction time variability [across trials standard deviation (SD)] (right). Black horizontal line depicts mean across participants and grey box ± standard error of the mean.

### Behavioural variability is increased in older adults

Reaction time variability was increased in older adults (Fig. 1B). The older participants were on average slower responding to the go stimulus, however this difference was not statistically significant [*t*_(73)_ = -1.84, *p* = .070]. Nevertheless, reaction time variability (within-subject across trials SD) was significantly higher in the older group [*t*_(73)_ = -2.53, *p* = .013]. As signal variability increases with signal mean, it is common to study the coefficient of variation (SD/mean) to estimate if the signal variability increased more than what would be expected from the increase in mean. Reaction time coefficient of variation was not significantly different across groups suggesting that the increase in reaction time variability was linked to the slowdown of the responses in the older group [*t*_(73)_ = -1.57, *p* = .120].

### Variability in the amplitude of preparatory evoked responses is decreased in the older group and predicts behaviour

Trial-by-trial variability in the CNV amplitude (CNV variability) was decreased in the older group (Fig. 2). During the preparatory period between the cue and the target, a frontal-central negative potential (the CNV) could be observed in the EEG of both groups of participants with strongest amplitude in the fronto-central electrode FCz (Fig. 2A and C). Although the amplitude of this evoked response was on average stronger (more negative) in the older group, trial-by-trial variability (across trials standard deviation) was reduced in the older group (Fig. 2B and D; see Suppl. Fig. 1 for single subject examples of trial-by-trial CNV variability). We observed significant group differences in the CNV amplitude and variability (Fig. 2E and F). However, the group differences showed different topographic distributions. CNV amplitude presented strongest group difference in left central electrodes, while CNV variability presented strongest group difference in parieto-occipital electrodes.

**Figure 2.**
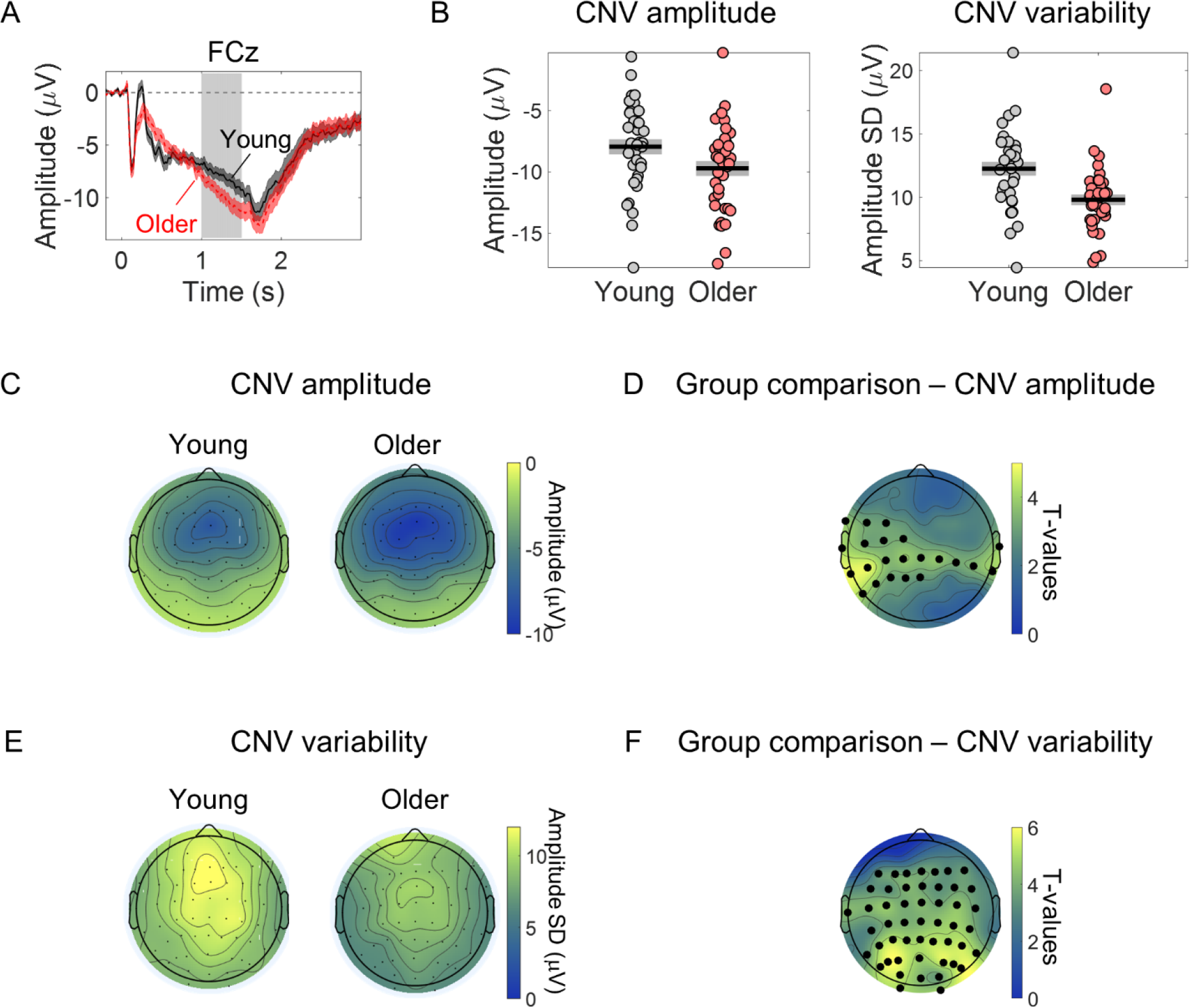
CNV amplitude and CNV variability show significant age-related differences. (A) Cue-locked ERP measured in the electrode FCz (mean ± SEM across participants). Grey background highlights time window used to average the ERP amplitude within each trial to study group differences in CNV amplitude and across trials CNV amplitude variability (1 – 1.5 s after cue onset). (B) Left, FCz CNV amplitude. Right, FCz CNV variability (across trials SD). Graphs depict individual data points (circles), mean (black horizontal line) and ± SEM across participants (grey box). (C) Scalp topography depicting average CNV amplitude. (E) Scalp topographies of CNV variability. (D and F) Head plots represent scalp topographies of *t*-values from independent samples *t*-tests comparing young and older participants at each electrode location. Correction for multiple comparisons was achieved using permutation tests and the ‘‘tmax’’ method for adjusting the *p* values (Blair & Karniski, 1993; Groppe et al., 2011). Black circles highlight the electrodes where the group difference was significant after controlling for multiple comparisons (*p* <.05).

Trial-by-trial CNV amplitude fluctuations predicted behaviour fluctuations (Fig. 3). Robust within-subject correlation analyses revealed a significant relationship between CNV amplitude and reaction time. Although, the correlation coefficients were on average smaller (closer to zero) in the young group, these did not differ significantly across groups (|*t*s| < 2.6). The correlation values were significantly different from zero in a left fronto-central cluster of electrodes (one-sample *t*-tests including all participants; Fig. 3A).

**Figure 3.**
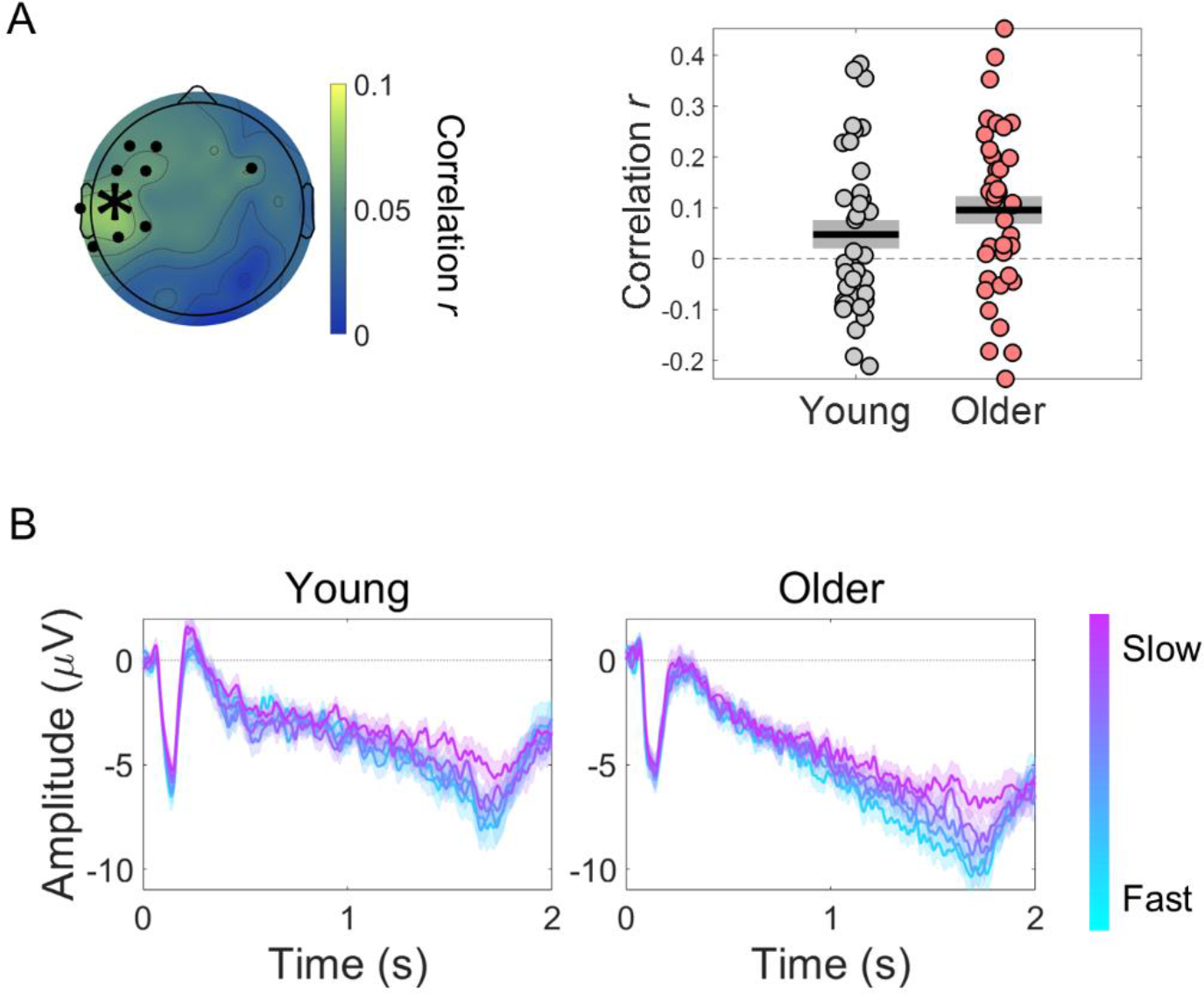
Within-subject correlation analyses between CNV amplitude and reaction time. (A) Scalp topographies of average correlation coefficients. Black circles highlight electrodes where correlation coefficients were significantly different from zero after controlling for multiple comparisons. Graph on the right shows individual data points (circles), mean (black horizontal line) and ± standard error of the mean across participants (grey box) of correlation coefficients for electrode C5 marked in the scalp topography with an asterisk. (B) ERPs at electrode C5 divided within each participant in five bins of trials according to reaction time. Fast responses were associated with more negative ERPs in both groups. Data are represented as mean ± standard error of the mean across participants.

Older individuals presented reduced trial-by-trial variability in cue-locked pupil dilation (PD) responses (Fig. 4). The warning cue elicited a pupil dilation response that peaked around 3 s after cue-onset (as reported before in Ribeiro and Castelo-Branco, 2019a; replotted in Fig 4A). PD response amplitude was not significantly different across groups [*t*_(71)_ = -1.82, *p* = .073]. The variability of PD response amplitude was, however, significantly reduced in the older group [*t*_(71)_ = 6.05, *p* < .001; Fig. 4B; see Suppl. Fig 1 for single subject examples of trial-by-trial variability in evoked responses].

**Figure 4.**
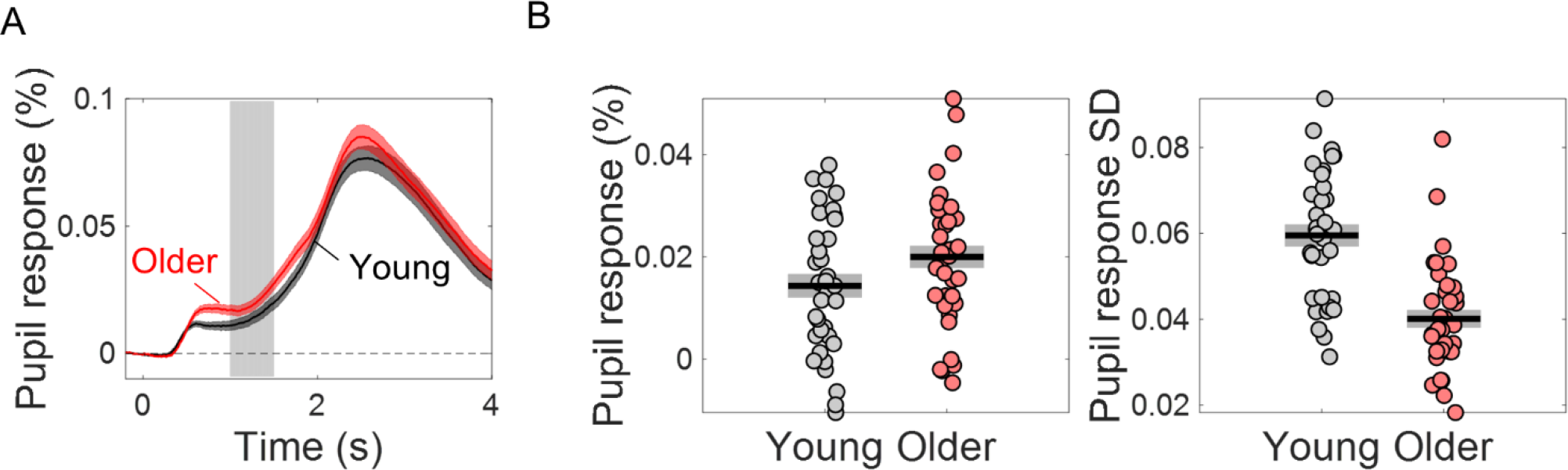
The variability of the cue-locked pupil dilation (PD) response is significantly reduced in older individuals. (**A**) Cue-locked pupil dilation response in young and older participants (mean ± SEM across participants). Grey background highlights time window (1 – 1.5 s after cue onset) where the PD amplitude was averaged within each trial to study the pupil dilation response elicited by the cue stimulus during the preparatory period before target onset. (**B**) Left, amplitude of PD response in young and older adults. Right, trial-by-trial variability in PD response (across trials standard deviation of PD amplitude) in young and older adults. Graphs depict individual data points (circles), mean (black horizontal line) and ± standard error of the mean across participants (grey box).

Preparatory pupil dilation responses did not correlate with reaction time. Within-subject robust correlation coefficients were not significantly different across groups [*t*_(71)_ = 1.09, *p* = .278] or significantly different from zero [one-sample *t*-test including all participants, *t*_(72)_ = .667, *p* = .507] In summary, trial-by-trial variability in the amplitude of the evoked responses was markedly reduced in older adults, while reaction time variability was increased. These observations suggest that the additional variability observed in the young group originates in a signal component that is not behaviourally relevant. In the following sections, we will delve into the origin of the variability in these evoked responses.

### Spectral properties of the ongoing EEG and pupil signals

Evoked neural responses occur on a background of ongoing neural activity that unavoidably adds to the responses measured (Fox et al., 2006; Shimaoka et al., 2019). It is possible therefore that age-dependent changes in the dynamics of ongoing activity explain the differences in the variability of task-related evoked responses. We explored the relationship between ongoing signal dynamics and the variability of the evoked responses by studying the spectral properties of ongoing EEG and pupil data and how these relate to the evoked responses.

The ongoing EEG signal of older adults presented power spectral density (PSD) data that are less steep, with reduced offset (reduced amplitude at low frequencies), reduced alpha power and increased beta power (Fig. 5). Ongoing EEG activity was estimated from the pre-stimulus period (3.5 s) just before cue onset, taking advantage of the long inter-trial intervals used. We analysed the PSDs using the FOOOF toolbox that separates the aperiod from the periodic component of the spectra (Donoghue et al., 2020). The aperiodic component can be described by its two parameters: the spectral exponent (a measure of how steep the spectrum is) and the offset (which reflects the uniform shift of power across frequencies). Both were significantly reduced in the older group (Fig. 5B). Occipital alpha power was reduced in older individuals mainly in occipital channels. In contrast, beta power in frontal and central EEG channels was increased in the older group.

**Figure 5.**
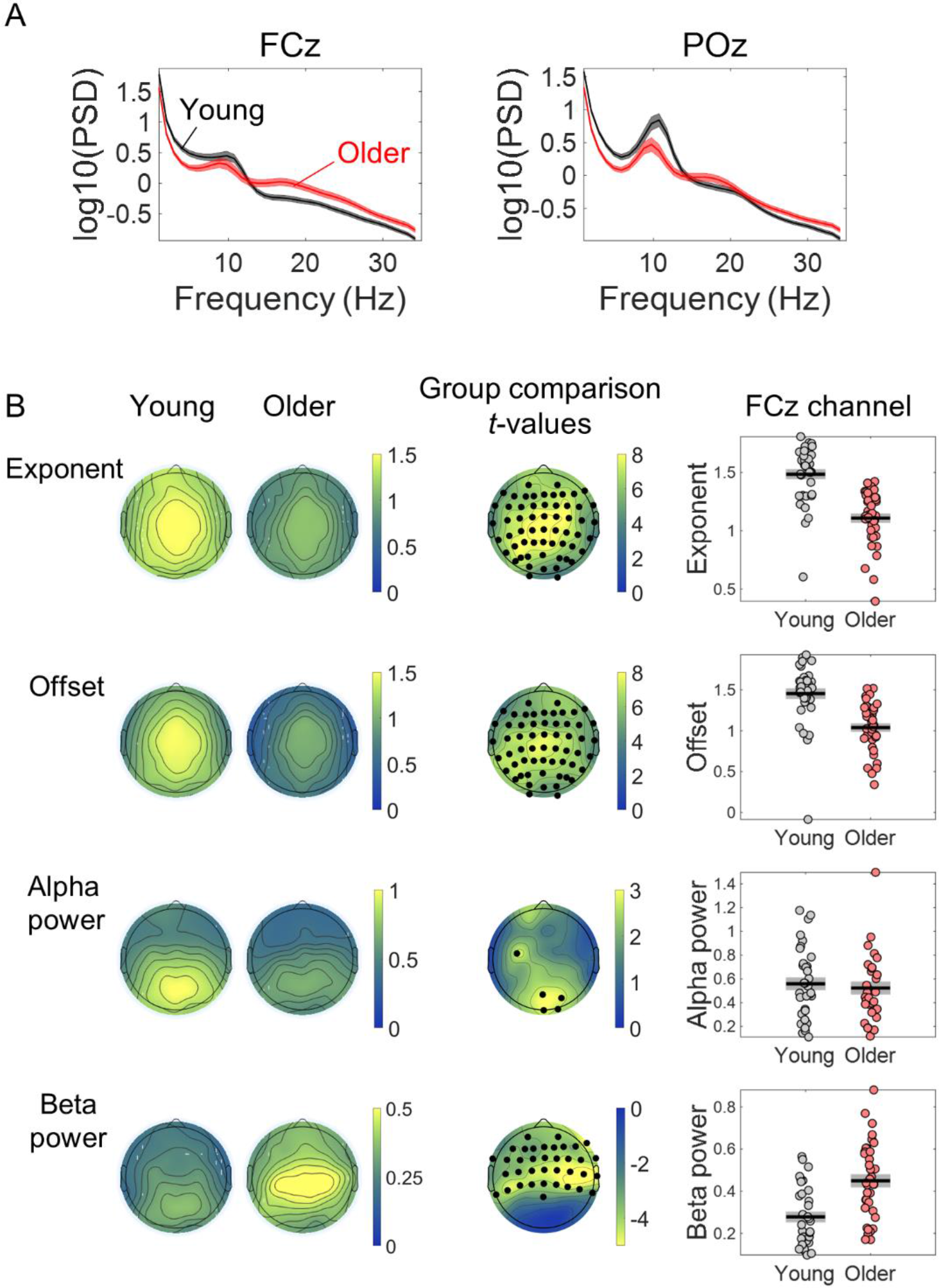
Comparison of the properties of the power spectral density (PSD) of the EEG pre-stimulus signal of young and older adults. (A) PSDs of the EEG signal in a window of 3.5 s before cue-onset measured in two EEG channels: the FCz, a frontocentral electrode where the preparatory CNV had maximal amplitude, and the POz, a parietocentral electrode where alpha oscillations were most prominent. Plots depict across participants mean ± standard error of the mean. (B) Scalp topographies of the average aperiodic parameters, exponent and offset, and alpha and beta power, extracted from the pre-stimulus PSD of each group of participants. Group differences for each of the parameters studied were estimated using independent *t*-tests. Channels that showed significant differences after controlling for multiple comparisons are highlighted in black. The graphs on the right show the exponent, offset, alpha power and beta power of the PSDs of young and older groups measured in the channel FCz. Graphs depict individual data points (circles), mean (black horizontal line) and ± standard error of the mean across participants (grey box). Participants where alpha or beta peaks were not detected were excluded from these graphs.

The PSDs of ongoing pupil signals of older individuals presented reduced exponent and offset (Fig. 6). Ongoing pupil signal fluctuations happen on a slower time scale than the EEG fluctuations, therefore, it is important to analyse longer time periods to study their dynamics. We took advantage of the pupillary recordings obtained at the beginning of the acquisition protocol while participants were fixating and passively listening to the cue stimulus being presented with the same frequency as in the task (Ribeiro & Castelo-Branco, 2019a). Due to the absence of task or luminance changes, the dynamics of pupil fluctuations in these recordings can be mostly assigned to ongoing pupillary activity. We calculated the pupil PSDs on 20 s epochs. The spectra of the ongoing pupillary signal followed a 1/f distribution, with power decreasing with increasing frequency (Fig. 6). The aperiodic parameters were significantly decreased in the older group [exponent: *t*_(70)_ = 7.48, *p* < .001; offset: *t*_(70)_ = 4.17, *p* < .001], i.e., the older group presented a flatter pupil spectrum with lower amplitude at the low frequencies. No obvious peaks were detected in the pupil spectra.

**Figure 6.**
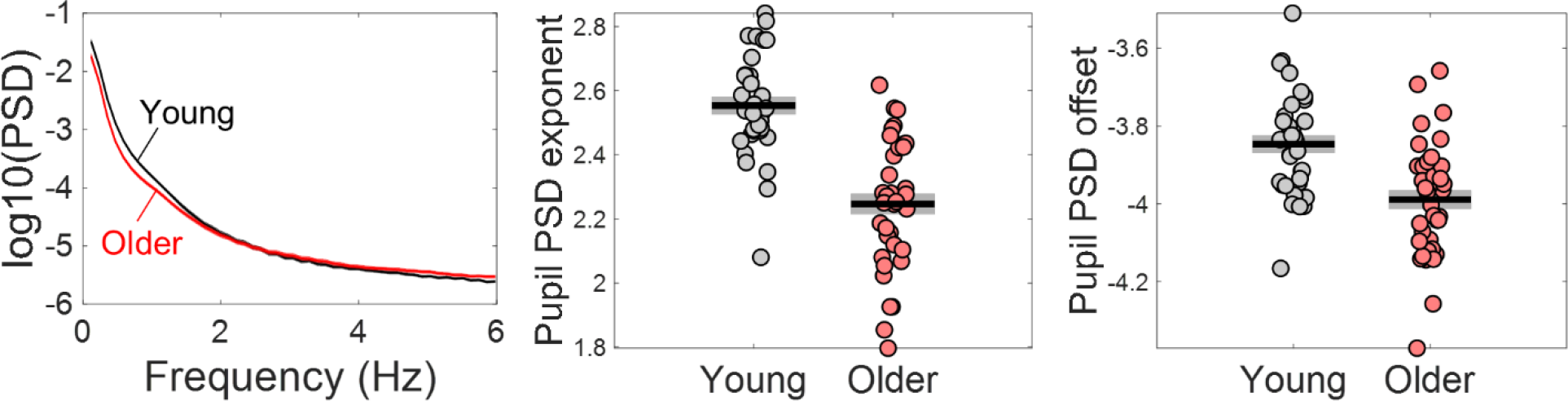
Power spectral density (PSD) of ongoing pupil signals acquired during passive fixation under constant luminance conditions. Left, PSD graph represents across participants mean ± standard error of the mean of the young and older group. Centre and right, graphic representation of the aperiodic parameters exponent and offset, respectively. Graphs depict individual data points (circles), mean (black horizontal line) and ± standard error of the mean across participants (grey box).

### Reduced variability in the evoked responses of older adults is linked to reduced amplitude in the slow fluctuations of ongoing activity

Spectral properties of ongoing signals predict trial-by-trial variability in evoked responses (Table 1). We used correlational analyses to investigate if the dynamics of the ongoing EEG and pupil signals captured by the spectral parameters was associated with the variability in task-related evoked responses across participants. For the EEG, we focused this analysis on the FCz electrode where the CNV showed its highest amplitude. Participants with higher spectral offset and exponent presented higher CNV and PD variability (Table 1; Supplementary Fig. 2). Alpha and beta power were not associated with CNV variability. These findings suggest that the aperiodic signal fluctuations are intrinsically linked to the variability observed in the evoked responses. We then sought to determine which frequencies in the aperiodic spectra were more predictive of the variability in the amplitude of the evoked responses. We found that the correlation between variability in the amplitude of the evoked responses and the amplitude of the power spectra was particularly strong in the lower frequencies (Supplementary Fig. 3). Importantly, the correlations were significant in both the aperiodic component of the fitted (FOOOF) spectra and in the raw (total) power spectra. In the total spectral power, the correlation between PD variability and spectral power was highest around .44 Hz and decreased for lower frequencies. In the EEG analyses, correlation was highest at 1 Hz, the lowest frequency analysed. These findings suggest that slow aperiodic fluctuations observed in the ongoing signals predict trial-by-trial variability in the amplitude of the evoked responses, both in the EEG and in the pupillogram.

**Table 1.** Pearson correlation between variability in the amplitude of the evoked responses and spectral parameters of the ongoing activity.

|  | Young | Older |
| --- | --- | --- |
| CNV SD |  |  |
| PSD exponent | **$r = .462$** $p = .005$ | $r = .136$ $p = .421$ |
| PSD offset | **$r = .673$** $p < .001$ | **$r = .538$** $p = .001$ |
| Alpha power | $r = .312$ $p = .068$ | $r = .168$ $p = .321$ |
| Beta power | $r = .160$ $p = .359$ | $r = -.004$ $p = .980$ |
| PD SD |  |  |
| PSD exponent | $r = .303$ $p = .077$ | $r = .182$ $p = .273$ |
| PSD offset | $r = .320$ $p = .061$ | **$r = .450$** $p = .005$ |

Significant correlations are displayed in bold font. SD = across trials standard deviation; PD = pupil dilation; CNV = contingent negative variation; PSD = power spectral density.

Adjusting for group differences in the power of the slow ongoing fluctuations eliminated the group differences in the variability of the evoked responses (Fig. 7). Once the effect of the spectral exponent, offset or slow spectral power were regressed out, the CNV variability was no longer significantly different across groups [adjusting for exponent *t*_(70)_ = .984, *p* = .328; adjusting for offset *t*_(70)_ = -.042, *p* = .966; adjusting for slow spectral power *t*_(70)_ = 1.20, *p* = .235; Fig. 7]. The group difference in the PD variability was also completely explained by the power of slow fluctuations observed in the ongoing pupillary signal [*t*-test group comparison no longer significant: *t*_(71)_ = 1.54, *p* = .129; Fig. 7]. However, adjusting for spectral exponent or offset did not completely explain the group difference [adjusting for exponent *t*_(71)_ = 2.45, *p* = .017; adjusting for offset *t*_(71)_ = 3.65, *p* < .001]. These findings indicate that the age-related reduction in the power of the slow aperiodic fluctuations underlie the group differences in trial-by-trial variability in the amplitude of the evoked responses.

**Figure 7.**
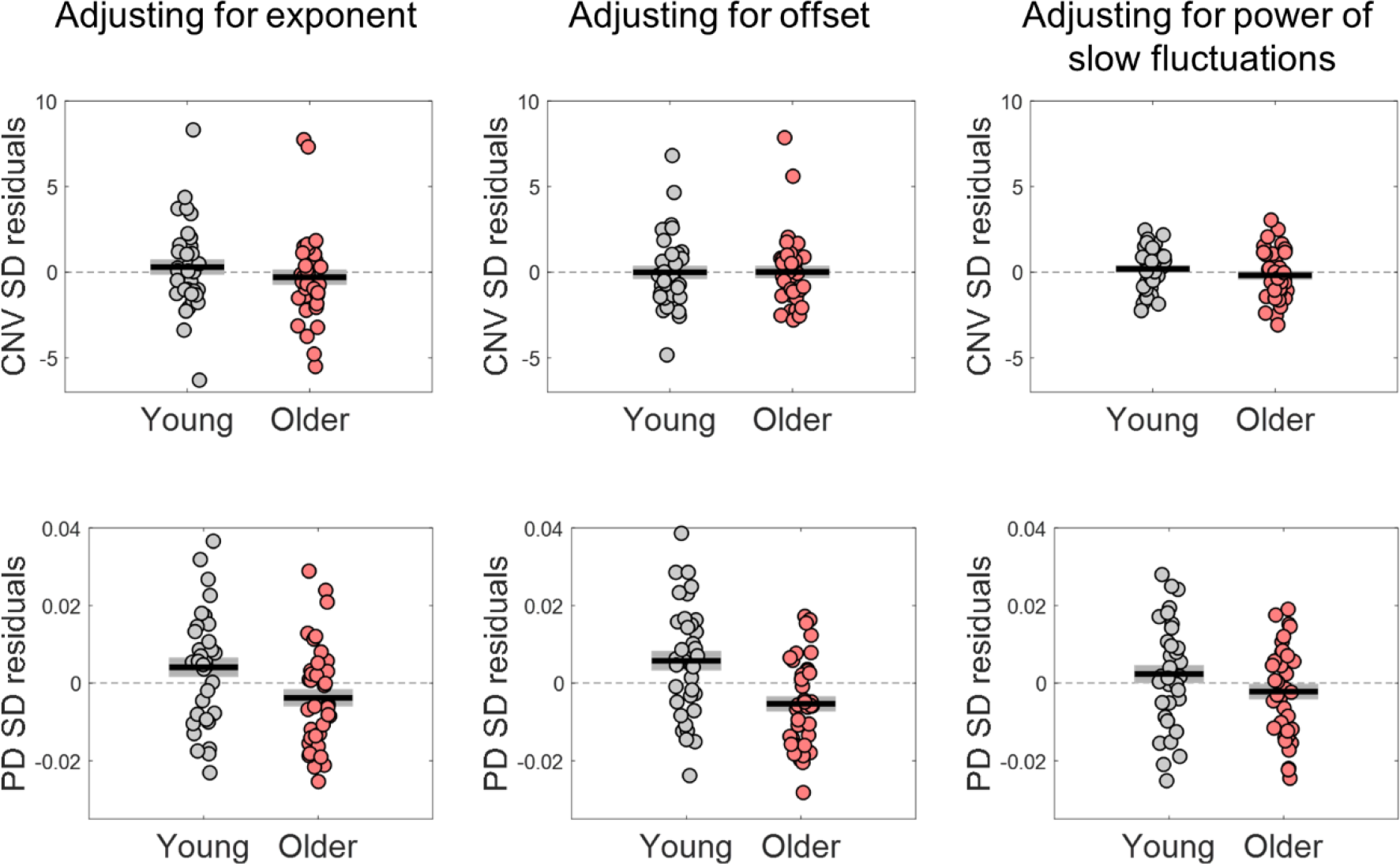
Variability in the amplitude of the evoked responses after adjusting for differences in the parameters of the power spectral density (PSD) of the respective ongoing signals. Top, variability in CNV amplitude measured at electrode FCz; bottom, variability in pupil dilation (PD) responses. Left, adjusted for the PSD’s exponent; centre, adjusted for the PSD’s offset; right, adjusted for the power of the slow fluctuations (power at 1 Hz in the EEG and power at .44 Hz in the pupil signal). Graphs depict individual data points (circles), mean (black horizontal line) and ± standard error of the mean across participants (grey box).

### Factors that contribute to trial-by-trial variability in evoked responses

There are two possible mechanisms through which background signal fluctuations affect the amplitude of the evoked responses. Background signal fluctuations might reflect fluctuations in brain state and different brain states will be associated with different evoked responses; and/or background signal fluctuations and the evoked responses are two independent signals that sum giving rise to a composite signal where the evoked response appears on top of a fluctuating baseline thereby modulating its shape and amplitude - a simple additive mechanism. In this latter case, the phase of the background fluctuations at the cue-onset rather than the instantaneous amplitude would better capture the relationship between the measured evoked responses and the ongoing signal (at the same amplitude values, the phase of the oscillation will define if the signal is increasing or decreasing). We explored this relationship by estimating the phase of the slow fluctuations in the EEG and pupil signals at cue-onset. For illustrative purposes, we plotted the relationship between the pre-stimulus phase and the amplitude of the respective evoked responses (Fig. 8). The evoked responses showed a dependence on the phase of the ongoing fluctuations in line with what would be expected if the measured evoked responses were the summation of an evoked response with an ongoing fluctuating signal (Fig. 8B and C). This relationship was also different from what would be expected if the evoked responses were affected solely by the amplitude of the baseline signal. This is further explored quantitatively in the next paragraph.

**Figure 8.**
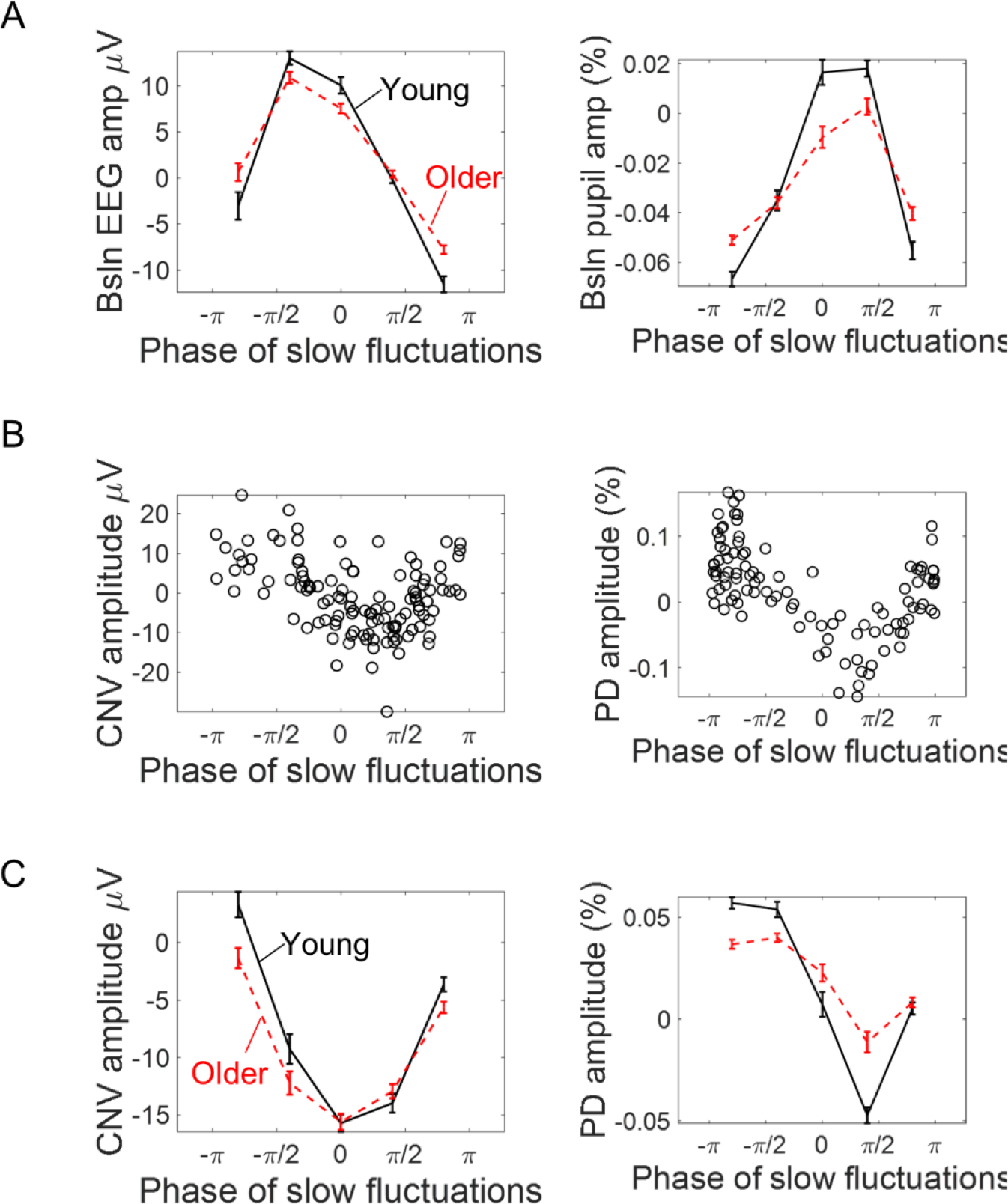
Association between the phase of slow ongoing fluctuations at cue-onset and the amplitude of the evoked responses in the EEG (left) and pupil (right) signals. (A) Baseline (pre-cue) signal amplitude sorted accordingly to the estimated signal phase (mean ± standard error of the mean across participants). As expected, signal amplitude reaches its peak at zero phase and its minimum at ± π. (B) Example data from two participants showing how the single trial amplitude of the evoked responses changed with the phase of the ongoing slow fluctuations. (C) Amplitude of the evoked responses split into five bins of trials (within each participant) according to the phase of the ongoing signals at cue-onset (mean ± standard error of the mean across participants).

Ongoing signal fluctuations strongly affected the amplitude of the evoked responses (Fig. 9). We quantified the association between EEG/pupil phase of slow fluctuations estimated at cue-onset (θ) and the amplitude of the respective evoked responses using linear regression. This relationship can be captured in a linear regression by representing the phase as a pair of variables: the cosine and the sine (Nurhab et al., 2017). We multiplied the cosine and sine functions by the amplitude envelop (r) of the slow signal fluctuations because the amplitude envelop varies throughout the recordings and these fluctuations will have differential effects on the evoked responses depending on its amplitude. The two predictors associated with the slow ongoing signal fluctuations were *r • sin θ* and *r •* cos θ. Note that *r •* cos θ is equal to the signal amplitude and therefore to the amplitude at baseline, while *r sin θ* will capture the additional effect of the signal phase. To study how these factors affected the trial-by-trial amplitude of the task-related responses, we run generalized linear regression models (GLMs) for each participant and separately for each signal type - EEG and pupil. We included as dependent variable the amplitude of the evoked response and as predictors *r • sin θ*, *r •* cos θ and, additionally, time-on-task (that can lead to differential responses due to learning, adaptation, or increased fatigue), and pre-stimulus alpha power (that reflects moment-to-moment spontaneous fluctuations in brain state related to alertness or attentional fluctuations; van Diepen et al., 2015). Run was included as categorical predictor (two consecutive runs of 8 minutes each were acquired). These models explained a large amount of the evoked responses variance (*R*_2_ mean ± SE: EEG young 67 ± 8 %; EEG older 63 ± 8 %; pupil young 73 ± 1 %; pupil older 67 ± 1 %). The explained variances were significantly decreased in the alternative model without the *r • sin θ* predictor that captured the effect of the ongoing signal phase (mean ± SE without *r • sin θ:* EEG young 27 ± 11 %; EEG older 28 ± 10 %; pupil young 40 ± 1 %; pupil older 37 ± 1 %, paired *t*-test comparing the two models’ explained variance: EEG *t*_(73)_ = -26.8, *p* < .001; pupil *t*_(71)_ = -22.9, *p* < .001), suggesting that the effect of the ongoing signal on the amplitude of the evoked responses is best captured by including amplitude and phase of the ongoing signal at baseline. We tested the estimated coefficients against zero using one-sample *t*-tests. The EEG GLM revealed that the CNV amplitude depended on time-on-task [it became less negative with time-on-task; *t*_(73)_ = 2.84, *p* = .006; Fig. 9A], and on the phase of the ongoing fluctuations at cue-onset captured by the cosine [*t*_(73)_ = -36.3, *p* < .001] and the sine [*t*_(73)_ = -81.7, *p* < .001] functions (Fig. 9A). It was independent of run [*t*_(73)_ = .035, *p* = .973] and pre-stimulus alpha power [*t*_(73)_ = -.530, *p* = .598]. Time-on-task had a stronger effect on the evoked responses of the young group in comparison with the older group [independent samples *t*-test: *t*_(72)_ = 3.74, *p* < .001]. The other coefficients did not present significant effects of group [run: *t*_(72)_ = 1.98, *p* = .051; pre-stimulus alpha: *t*_(72)_ = -.098, *p* = .923; cosine: *t*_(72)_ = .189, *p* = .851; sine: *t*_(72)_ = -.007, *p* = .994]. The pupil GLM revealed that the only predictors that significantly contributed towards trial-by-trial PD variability were the cosine and the sine of the pre-stimulus phase [one-sample *t*-test testing estimated coefficients against zero - cosine: *t*_(71)_ = -38.2, *p* < .001; sine: *t*_(71)_ = -51.3, *p* < .001]. The other predictors were not significantly different from zero [run: *t*_(71)_ = 1.88, *p* = .064; time-on-task: *t*_(71)_ = -1.34, *p* = .185; alpha power: *t*_(71)_ = - .061, *p* = .952]. None of the coefficients showed a significant group difference [run: *t*_(70)_ = -.198, *p* = .844; time-on-task: *t*_(71)_ = -1.00, *p* = .321; alpha power: *t*_(70)_ = 1.68, *p* = .098; cosine: *t*_(70)_ = -.143, *p* = .887; sine: *t*_(70)_ = -.396, *p* = .693]. These results suggest a strong effect of the amplitude and phase of the ongoing signal fluctuations on the evoked responses (Fig. 9A and B), consistent with the idea that there is a drift in the signal baseline that adds to the evoked responses pushing them towards positive or negative values depending on the phase at cue-onset.

**Figure 9.**
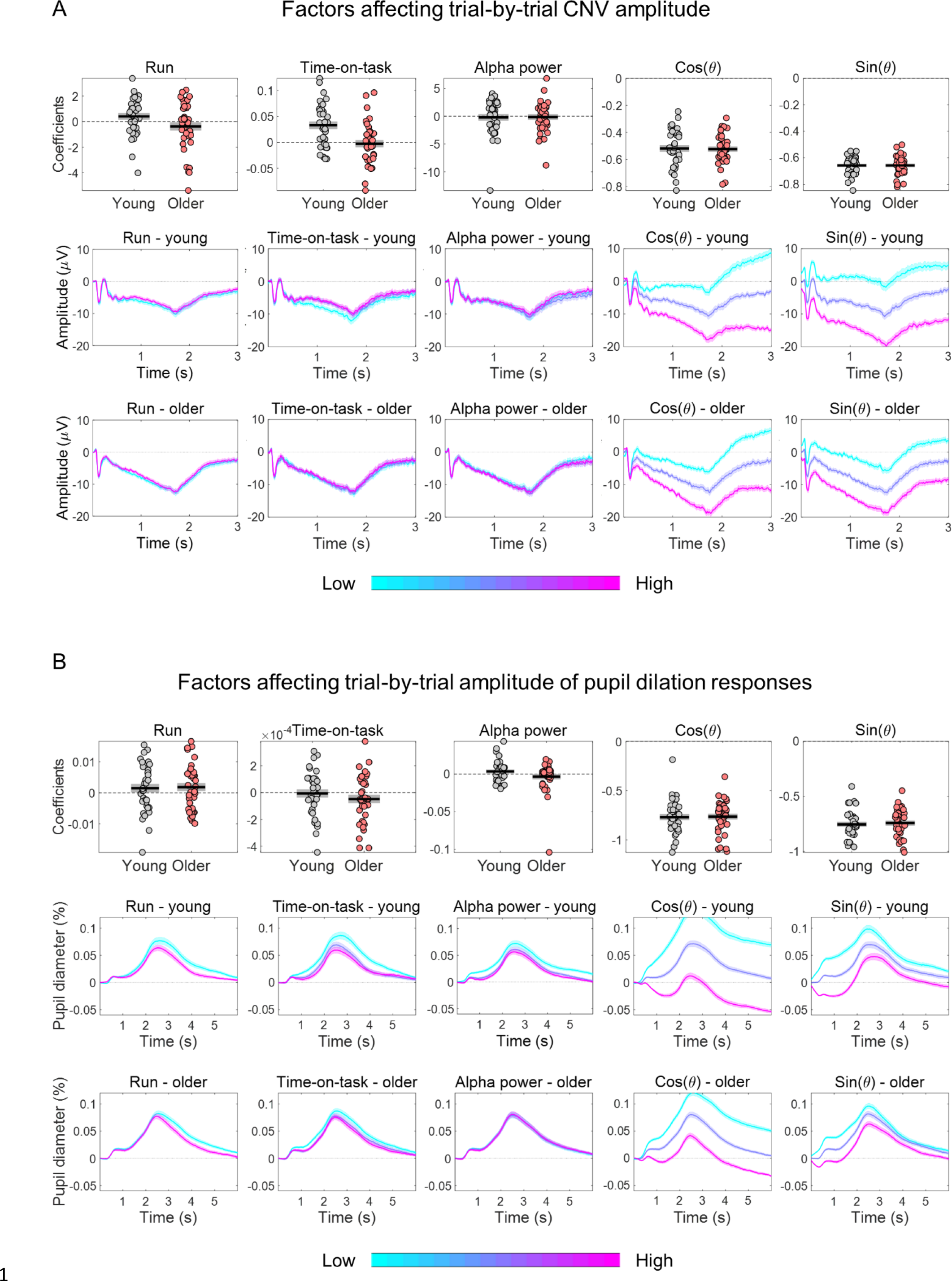
Estimated coefficients from within-subject generalized linear regression models (GLMs) including as independent variable the amplitude of the evoked response and as predictors the effects of time-on-task, pre-stimulus alpha power and the cosine and sine of the phase at cue-onset of the ongoing slow fluctuations. Run was included as categorical predictor. (A) Effect of predictors on CNV amplitude measured at FCz. (B) Effect of predictors on pupil dilation response (PD) amplitude. (A and B) Top, graphs depicting estimated coefficients (circles = individual data points; black horizontal line = across participants mean; grey box = ± standard error of the mean across participants). Middle and bottom, cue-locked evoked responses divided in three bins of trials (within each participant) sorted according to each of the predictors (mean ± standard error of the mean across participants). Note, that the graphs show raw responses not adjusted for the other predictors.

To test if the variability in the evoked responses associated with the ongoing signal fluctuations was behaviourally relevant, we studied how adjusting for the signal phase at baseline changed the correlation between the evoked responses and reaction time. After regressing out the effect of the ongoing signal fluctuations from the amplitude of the evoked responses, these predicted reaction time at least as strongly as without the adjustment. In particular, after regressing out the effect of the phase of the slow fluctuations at baseline, the CNV amplitude residuals correlated with reaction time in a larger number of electrodes than without the adjustment and unmasked a significant relationship between the CNV amplitude and reaction time in frontal electrodes that was not observed without the adjustment (Fig. 10A), suggesting that the relationship between CNV amplitude and behaviour is independent of the baseline fluctuations. On average the correlation coefficients with the adjustment were slightly more positive than without the adjustment. However, this difference was only significantly different in one EEG channel (F4) after correcting for multiple comparisons. For the pupil responses, we found that regressing out the effect of ongoing signal fluctuations was crucial to detect the link between PD evoked responses and reaction time. Indeed, after adjusting the PD amplitude for the effect of the phase of the signal fluctuations at baseline, the PD amplitude residuals correlated with reaction time [Fig. 10C; one-sample *t*-test: *t*_(72)_ = -9.55, *p* < .001]. Notably, the correlation coefficients with the adjustment for the phase of the ongoing signal were significantly different from the coefficients without the adjustment [paired *t*-test comparing with and without adjustment: *t*_(72)_ = -7.54, *p* < .001]. These findings suggest that the effect of the ongoing signal on the evoked responses is not behaviourally relevant and can mask brain-behaviour associations.

**Figure 10.**
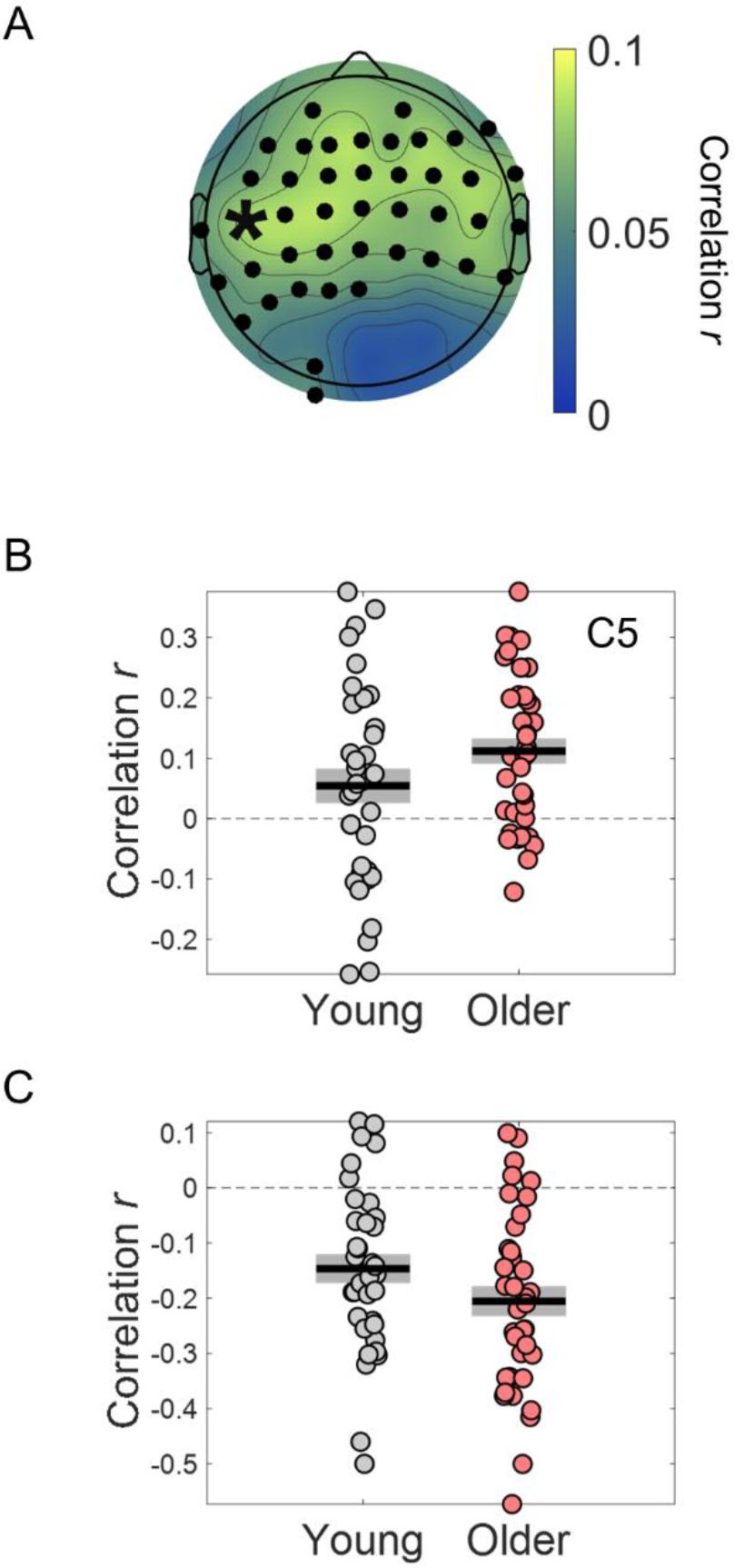
Within-subject robust Pearson’s correlation between single trial amplitude of the preparatory evoked responses adjusted for the effect of the ongoing signal fluctuations and reaction time. (**A**) Scalp topographies of robust Pearson’s correlation coefficients. Electrodes where the correlation coefficients were significantly different from zero are highlighted in black. (**B**) Example of correlation coefficients between the ERP amplitude adjusted for ongoing signal phase fluctuations measured in the channel C5 (marked with an asterisk in A) and reaction time. (**C**) Robust Pearson’s correlation coefficients between the amplitude of the pupil dilation responses adjusted for the phase of the ongoing signal fluctuations and reaction time (circles = individual data points; black horizontal line = across participants mean; grey box = ± standard error of the mean across participants).

It is important to note that after regressing out the effect of the ongoing signal’s phase at cue-onset, the variability of the evoked responses was still significantly reduced in the older group (Supplementary Fig. 4), suggesting that the baseline (pre-stimulus) values are not enough to completely predict the effect of the ongoing signal fluctuations on the evoked responses in line with the idea that these slow signal fluctuations are aperiodic and therefore not completely predictable (Shimaoka et al., 2019).

## Discussion

Task-related brain evoked responses occur on top of ongoing signal fluctuations that markedly affect their amplitude on a trial-by-trial basis. Our results revealed that as a consequence of older people showing reduced amplitude in the ongoing slow signal fluctuations measured in the EEG and pupil signals, trial-by-trial variability in their evoked responses is also decreased. Once the difference in ongoing signal dynamics was adjusted for, the evoked responses were equally variable in young and older adults. This finding is consistent with the fact that behavioural variability does not decrease with age as would be expected if the evoked responses were more consistent across trials. In fact, although ongoing signal fluctuations strongly affected the amplitude of the evoked responses, these amplitude modulations did not appear to have behavioural consequences.

There are two possible ways through which the slow fluctuations observed in the ongoing signals impact the evoked responses. One possibility is that these slow signal fluctuations reflect fluctuations in brain state that affect the way the brain responds to external stimuli as has been suggested for example for the impact of infra-slow oscillations observed in EEG (Monto et al., 2008) or pre-stimulus activity levels in resting-state networks in fMRI (Boly et al., 2007). The other possibility assumes that the effect of the ongoing fluctuations on the evoked responses is purely additive and simply reflects the fluctuating baseline and is therefore not relevant for the type of task studied. This second hypothesis is supported by previous fMRI studies in humans (Fox et al., 2006; Schölvinck et al., 2012) as well as studies of neuronal responses measured in the sensory cortices of mice and cats (Scholvinck et al., 2015; Shimaoka et al., 2019). Our results further suggest that this latter case dominates the effect of the ongoing EEG and pupil signals on their respective evoked responses, highlighting an effect of ongoing signal’s phase: if the ongoing signal is increasing in amplitude, then the evoked response shows a more positive amplitude and vice-versa. Indeed, similar baseline amplitude levels at different signal phases (with amplitude increasing or amplitude decreasing) have different effects on the amplitude of the evoked responses (see Fig. 8) and the variance of the evoked responses explained by the models tested increased substantially by including the ongoing signal phase [r • sin(θ)] at cue-onset in comparison with including only the signal amplitude (as captured by the cosine function). Importantly, when we adjusted for the effect of the ongoing fluctuations on the evoked response’s amplitude, the correlation between the evoked response and reaction time was at least as strong as without this adjustment, suggesting that the effect of the ongoing signal fluctuations on the evoked responses has no impact on behaviour. Interestingly, in both the pupil and EEG signals adjusting for ongoing signal fluctuations unmasked a relationship between the evoked responses amplitude and reaction time that, without the adjustment, was not detectable in the pupil and was evident only in a restricted number of channels in the EEG. This observation suggests that the additional variability imposed on the evoked responses by the ongoing signal can mask brain-behaviour associations and therefore should be attended to and adjusted for.

Previous fMRI studies suggest that task evoked responses are affected by ongoing signal fluctuations occurring in the same brain areas but that these ongoing fluctuations are shared across brain regions (Fox et al., 2006; Schölvinck et al., 2012). Similarly, single cell electrophysiology and optical imaging with voltage sensors in anesthetized cats suggest that sensory responses in individual neurons are variable from trial-to-trial and that this variability reflects the sum of the sensory response with an ongoing signal shared across neurons (Arieli et al., 1996; Scholvinck et al., 2015). The CNV preparatory potential originates not from an unitary neural mechanism but from a set of brain areas, including the supplementary motor area (SMA), anterior cingulate cortex (ACC) and the thalamus (Nagai et al., 2004). Likely, these areas will display ongoing activity that will sum to the evoked responses. Nevertheless, the ongoing EEG fluctuations studied probably originate in a large set of brain areas that will include, but not be limited to, the brain areas involved in the preparatory mechanisms linked to the CNV potential.

Fluctuations in pupil size under constant illumination conditions reflect activity in a restricted number of brain regions. The main areas that have been identified in non-luminance modulation of pupil size are the superior colliculus and the locus coeruleus (LC; Joshi and Gold, 2020). The involvement of the superior colliculus in pupil dilation responses is thought to predominate during the orienting response to sensory stimuli (Wang et al., 2014), while activation of the LC modulates pupil size during resting periods as well as during cognitive arousal (Joshi et al., 2016; Murphy et al., 2014; Reimer et al., 2016). However, the pupil inverse problem (which brain areas are being modulated when we observe changes in pupil size) remains to be solved (Joshi & Gold, 2020). It is possible that, in warned reaction time tasks like the one used in the current study, ongoing fluctuations in pupil size reflect activity in a different brain area from the brain area eliciting the task-related responses. The idea that the ongoing pupil signal and the task-related pupil evoked response can be two largely independent signals, challenges previous beliefs. The prevailing model suggests that high baseline (tonic) activity in the LC is associated with weaker task-related (phasic) responses, and that pupil data parallels this observation (Aston-Jones & Cohen, 2005; Gilzenrat et al., 2010). However, the application of this model to human pupil data acquired during performance of typical psychophysical tasks has been challenged given the relatively small modulations in arousal that occur under these conditions (Joshi & Gold, 2020). Our analyses further suggest that the effect of pupil baseline on the pupil dilation responses is not behaviourally relevant and merely additive.

Our analyses showed that ageing affects the dynamics of the ongoing EEG signal (reflecting cortical activity) and the ongoing pupillary signal (reflecting activity in brainstem neuromodulatory systems). In both signals, older people presented flatter spectra with reduced power in the slower aperiodic fluctuations. This flattening of brain signal spectra with ageing has been described before in EEG and ECoG data (Voytek et al., 2015; Waschke et al., 2017), is also evident in MEG data (Vlahou et al., 2015), and is here reported in the pupil signal suggesting an age-related decrease in the amplitude of the slow fluctuations in cortical and subcortical networks. In line with our findings, Vlahou et al (2015) showed that, in MEG data, the power of slow fluctuations decreases linearly with ageing (Vlahou et al., 2015). Sequential activation and de-activation of resting-state networks is observed in the ongoing EEG data both during rest and task performance (Abreu et al., 2020; Liu et al., 2017). This ongoing dynamic possibly underlies the ongoing signal fluctuations we observe at the channel level EEG. The fact that EEG, MEG and pupil spectra decrease in amplitude with increasing frequency suggests that brain activation patterns that involve large activation volumes accumulate relatively slowly (the larger the brain volume recruited the slower the activity changes). Aging is characterized by a significant decrease in the strength of long-range structural connections (Westlye et al., 2010) as well as a decrease in the functional connectivity (correlated BOLD activation patterns) within large-scale brain systems, like the default mode network (Sala-Llonch et al., 2015), consistent with the idea that simultaneous activation of distant brain areas supported by long-range structural connectivity is impaired in older individuals (Andrews-Hanna et al., 2007). Reduced amplitude of slow EEG fluctuations might therefore be related to impaired recruitment of large-scale brain systems.

### Conclusions

The current study shows that the amplitude of the preparatory evoked potential, the CNV, and the preparatory pupil dilation responses show comparable trial-by-trial variability in young and older adults once the effect of the ongoing signal fluctuations is considered. This finding suggests a dissociation between ongoing brain activity, which presents reduced variability in older adults, and task-related responses, which does not. This dissociation might explain why although ongoing brain activity is less variable in older adults, behaviour variability increases with ageing.

## Materials and methods

### Participants

Thirty-six young adults (mean age ± SD = 23 ± 3 years; 29 women; 3 left-handed) and thirty-nine older adults (mean age ± SD = 60 ± 5 years; 31 women; 3 left-handed) were included in this study. Participants’ characteristics were reported elsewhere (Ribeiro & Castelo-Branco, 2019a). EEG data from one older participant and pupil data from one young adult and one older adult with light-coloured eyes were not included due to poor data quality.

The study was conducted in accordance with the tenets of the Declaration of Helsinki and was approved by the Ethics Committee of the Faculty of Medicine of the University of Coimbra. Written informed consent was obtained from the participants, after explanation of the nature and possible consequences of the study.

### Task design

The task was designed and run with the Psychophysics Toolbox, version 3 (Brainard, 1997), for Matlab (The MathWorks Company Ltd). Task design is schematized in Figure 1 and has been described in detail elsewhere (Ribeiro & Castelo-Branco, 2019a, 2019b). Briefly, we applied two cued auditory tasks: a cued simple RT task and a cued go/no-go task. In this study, for the sake of conciseness, we report only the analyses of the go/no-go task data. Equivalent results were obtained with the data from the simple RT task with the exception that PD amplitude, without the adjustment for the slow ongoing fluctuations, correlated with reaction time. Nevertheless, after the adjustment the correlation was significantly stronger. The auditory stimuli used were three different pure tones, suprathreshold, with duration of 250 ms, with the following frequencies: cue - 1500 Hz; go stimulus - 1700 Hz; no-go stimulus - 1300 Hz. Participants performed the two tasks, cued simple RT and cued go/no-go, sequentially. The order of the tasks was counterbalanced across participants, i.e., half of the participants started with the simple RT and the other half with the go/no-go. In the simple RT task, the cue was followed by the go stimulus (100 trials) to which participants were instructed to respond by pressing a keyboard key as fast as possible with their right index finger. In the go/no-go task, the cue was followed or by the go stimulus (80 trials) or by the no-go stimulus (20 trials). Participants were instructed to respond as fast as possible to the go stimulus with their right index finger, while refraining from responding to the no-go stimulus. The intertrial interval was variable with a median of 7.6 s (min 6.7 and max 19.6 s). The interval between the cue and the target stimuli and between target and the beginning of the next trial were drawn from a nonaging distribution, –W*ln(P), where W is the mean value of the interval distribution and P is a random number between 0 and 1 (Jennings et al., 1998). In our task design, the cue-target interval was 1.5-0.25*ln(P) and the interval between the target and the beginning of the next trial (cue) was 5.2-1*ln(P) in seconds.

In the analysis of task performance, we assessed reaction time and task accuracy. We considered as error trials all trials where the participants responded after cue presentation, failed to respond to the go stimulus (misses), responded to the go stimulus too slowly (slower than 700 ms), or responded to the no-go stimulus in the go/no-go condition. These trials were signalled with a feedback tone warning the participants that an error was committed.

### EEG data acquisition and pre-processing

As previously described (Ribeiro & Castelo-Branco, 2019a, 2019b), the EEG signal was recorded using a 64-channel Neuroscan system (Compumedics EUROPE GmbH) with scalp electrodes placed according to the International 10–20 electrode placement standard, with reference between the electrodes CPz and Cz and ground between FPz and Fz. Acquisition rate was 500 Hz. Vertical and horizontal electrooculograms were recorded to monitor eye movements and blinks. The participants’ head was stabilized with a chin and forehead rest to record pupillographic data simultaneously. Consequently, the electrodes on the forehead, FP1, FPz, and FP2, displayed signal fluctuation artefacts due to the pressure on the forehead rest. These were excluded from the analyses. A trigger pulse was generated at the onset of each stimulus and at every button press. EEG data analysis was performed with the EEGLAB toolbox versions 14.1.1 and 19.1 (Delorme & Makeig, 2004) and Matlab custom scripts.

We used independent component analysis (ICA) to eliminate nonbrain artefacts from the data, as described previously. The data, re-referenced to linked earlobes, was then band pass filtered with cut off frequencies of 0.1 and 35 Hz, and periods containing further artefacts were manually removed.

### Pupil data acquisition and pre-processing

As described previously (Ribeiro & Castelo-Branco, 2019a), the pupil diameter of the right eye was measured by an infrared eye-tracker (iView X Hi-Speed 1250 system from SMI) with a sampling rate of 240 Hz. Analysis of pupil data was performed using Matlab custom scripts and the EEGLAB toolbox in Matlab. Artefacts and blinks were corrected using the blink correction Matlab script by Greg Siegle (stublinks.m) available in http://www.pitt.edu/~gsiegle/ (Siegle et al., 2003). Briefly, artefacts, including blinks, were identified as large changes in pupil dilation occurring too rapidly to signify actual dilation or contraction. Linear interpolations replaced artefacts throughout the data sets. Data were smoothed using a 3-point unweighted average filter applied twice.

Pupil epochs were visually inspected for artefacts not adequately corrected by the linear interpolation procedure. Epochs with remaining artefacts were manually rejected. In addition, we discarded all error trials and correct trials that immediately followed error trials. The numbers of included trials did not differ significantly across groups for each condition.

For each run, pupil data was expressed as the percentage of the mean across the whole run by dividing the run data by the mean pupil size within that run.

### Single trial amplitude and variability of evoked responses

The preparatory evoked responses in the EEG and pupil signals were studied in cue-locked epochs with average baseline amplitude (200 ms time window before cue onset) subtracted. Single trial amplitude was calculated by averaging the cue-locked responses within the time window from 1 s to 1.5 s after cue-onset. This time window was positioned just before the earliest target onset.

Group comparison of average CNV amplitude and across trials SD of CNV amplitude was performed, for each electrode, with independent samples permutation test based on a *t*-statistic. The “tmax” method was used for adjusting the *p* values of each variable for multiple comparisons, across the 59 electrodes analysed (Blair & Karniski, 1993; Groppe, 2021b; Groppe et al., 2011).

### Correlation between single trial amplitude of evoked responses and reaction time

For within-subject correlation analyses between reaction time and single trial amplitude of the evoked responses, we used the Pearson’s skipped robust correlation (Wilcox, 2012), as implemented in the robust correlation Matlab toolbox (Pernet et al., 2013; Rousseeuw, 1984). Skipped correlations minimize the effects of bivariate outliers by considering the overall structure of the data. Notably, Pearson’s skipped correlation is a direct reflection of Pearson’s *r*. At the group level, correlations were considered significant if the correlation coefficients were significantly different from zero. For the EEG, this was established using one sample permutation test based on a *t*-statistic with the “tmax” method used for adjusting the *p*-values of each variable for multiple comparisons (across the 59 electrodes analysed) (Groppe, 2021a; Groppe et al., 2011).

### Spectral analyses of ongoing signals

EEG and pupil power spectral densities (PSDs) estimated via the Welch’s method were fit using the FOOOF algorithm (version 1.0.0) using the Matlab wrapper (Donoghue et al., 2020).

In the EEG analyses, PSDs were estimated in single trials in 3.5 s long pre-cue epochs. PSDs were then averaged across trials for each electrode and fit using the FOOOF algorithm with the following settings: peak width limits = [1 8]; peak threshold = 1; aperiodic mode = ’fixed’; minimum peak height = 0.1; max number of peaks = 4; frequency range = [1 35]. Peaks with peak frequency between 7 and 14 Hz were considered within the alpha range and peaks with peak frequency between 14 and 30 Hz were considered within the beta range. Peaks outside these frequencies were rarely detected and were not analysed.

In the pupil analyses, PSDs were estimated in 20 s long epochs from a 4 min recording acquired at the beginning of the experiment during which the participants were passively fixating and listening to the cue stimulus being presented with the same frequency as in the cued auditory tasks but without any overt task (Ribeiro & Castelo-Branco, 2019a). The 20 s epochs were cut sequentially and were not locked to the auditory stimulus. These long epochs allowed us to assess the slow signal fluctuations typical of the pupillary recordings. Pupil PSDs were then averaged across trials and were fit using the FOOOF algorithm with the following settings: peak width limits = [-inf .25]; peak threshold = 3.5; aperiodic mode = ’fixed’; minimum peak height = 0.2; maximum number of peaks = 4; frequency range = [0.05 6].

FOOOF model fitting quality was estimated from the model *R*^2^. For the EEG data, average *R*^2^ across electrodes and across participants was .99 for the young group and .97 for the older group. These *R*^2^ values reflected good model fittings, however, it is important to note that they showed significant differences across groups in most EEG electrodes (Supplementary Figure 5A and B). As the *R*^2^ values were lower in the older group, it is possible that the parameter estimation was not as good in this group. Surprisingly, the model *R*^2^ correlated with the estimated values for exponent and offset – lower *R*^2^ values were associated with lower exponents and offsets (Supplementary Figure 5C). This relationship was evident in both groups of participants. To ensure that the study conclusions were not driven by differences in fitting quality, we compared the correlation between the variability in the evoked responses and the fitted spectra with the correlation between the variability in the evoked responses and the raw PSD (supplementary Fig. 3). We obtained similar results: the power of low frequencies in the aperiodic fitted spectra and in the raw spectra correlated with the evoked responses amplitude variability. The *R*^2^ for the fitting of pupil PSDs were not significantly different across groups (*R*^2^ young = .99; *R*^2^ older = .98; Supplementary Figure 5D). Pupil *R*^2^ correlated with the estimated PSD offset values but not with the PSD exponent.

For the correlation analyses between the spectral parameters of the ongoing signal measured at FCz and CNV variability, participants with spectral goodness-of-fit more than 2.5 standard deviations away from the mean, calculated as the z-score of *R* transformed into Fisher’s Z, were excluded from the analyses (one young and one older adults). These were also the participants with lowest spectral exponent.

### Estimation of the phase of the slow fluctuations in the EEG and pupil signals

The instantaneous phase and amplitude envelope of the slow fluctuations was estimated using the Hilbert transform. Before applying the Hilbert transform, the pre-processed EEG signal was bandpass filtered between .1 Hz and 2 Hz and the pre-processed pupil signal was bandpass filtered between .1 and .9 Hz. Phase (θ) and amplitude envelope (r) at the time of cue-onset were extracted and used to calculate two variables that represent the circular phase of the signal in Cartesian coordinates: *r • sin θ* and *r •* cos θ (Nurhab et al., 2017). These variables were included as predictors in the linear regressions studying the effect of pre-stimulus phase angle on the amplitude of the evoked responses. Note that *r •* cos θ is equal to the amplitude of the filtered signal at each time point.

### The effect of pre-stimulus variables on single trial amplitude of evoked responses

We run within-subject generalized linear regression models, with the single-trial amplitude of the cue-locked evoked responses (average within the time interval from 1 to 1.5 s after cue-onset) as response variable and including run as categorical predictor (two sequential runs were acquired per participant), and, as continuous predictors, time-on-task (defined as the trial number within each run), alpha power (estimated from the FOOOF model fitting of 3.5 s pre-stimulus epochs in electrode POz where it showed the highest amplitude; when no alpha peak was detected, alpha power was set at zero), r • sin θ and r • cos θ (estimated as described above for each signal).

The models explained a large amount of variance of the evoked responses amplitude, however, the explained variance was higher in the young than in the older group [independent samples *t*-test comparing the models’*R*^2^ of both groups: EEG *t*_(72)_ = 2.11 *p* = .039; pupil *t*_(70)_ = 2.88, *p* = .005].

### Adjusting single trial amplitude of the evoked responses for pre-stimulus phase angle of ongoing signals

To adjust the evoked responses amplitude for the effect of ongoing signal fluctuations, we regressed out the effects of the phase of the signal at cue-onset by taking, for each participant, the residuals from the multiple linear regression with response variable the evoked responses amplitude and r • sin θ and r • cos θ as predictors. The regression was run separately for each signal (EEG and pupil).

## Acknowledgements

This work was supported by Fundação para a Ciência e a Tecnologia (grants references: FCT/UIDB&P/4950/2020, PTDC/PSIGER/30852/2017, DSAIPA/DS/0041/2020).

## Data availability

Data has been deposited to Open Neuro Repository under the doi: 10.18112/openneuro.ds003690.v1.0.0

## Competing interests

Declarations of interest: none.

## SUPPLEMENTARY MATERIAL

**Supplementary Figure 1.**
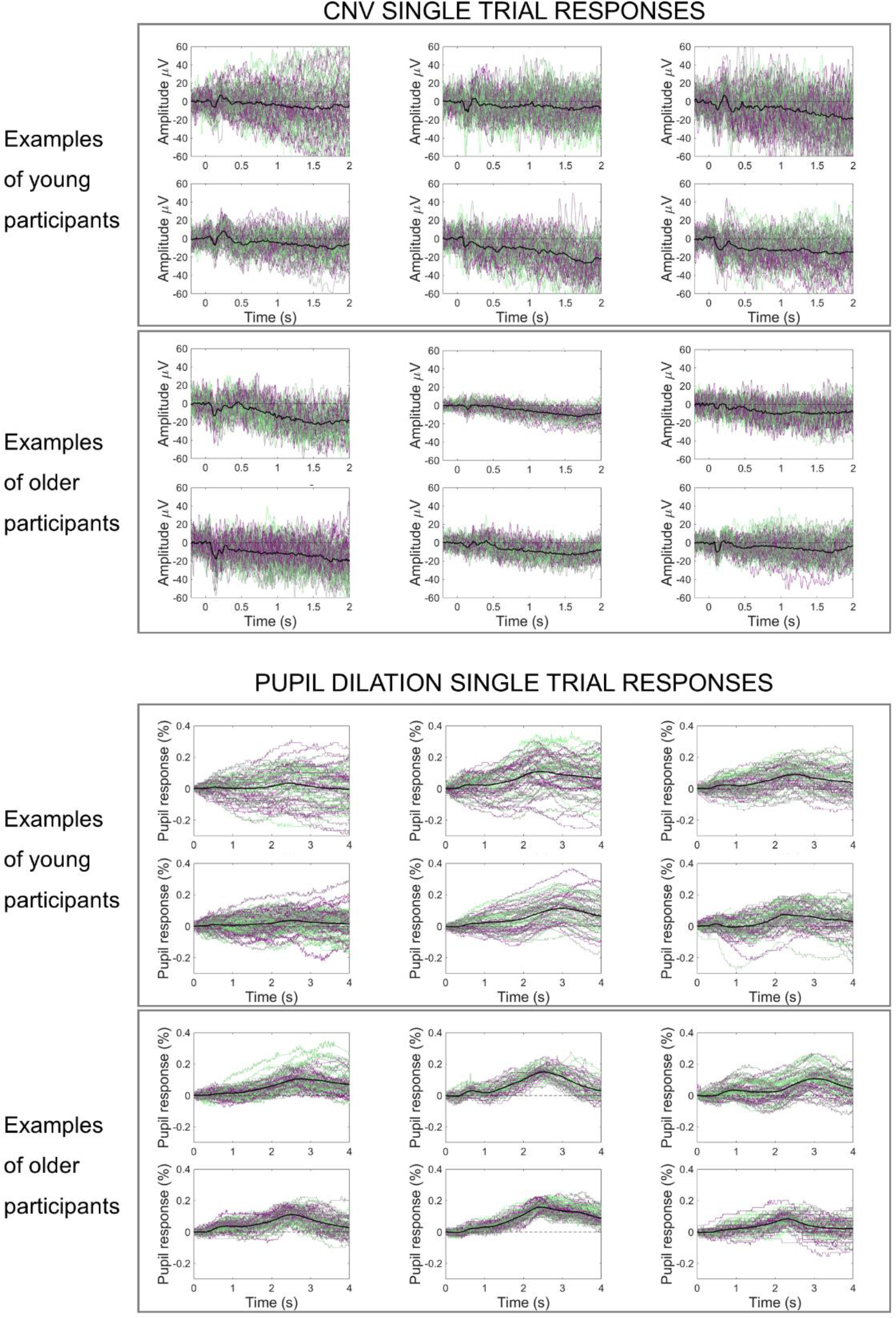
Examples of cue-locked evoked responses from young and older participants showing higher trial-by-trial variability in the evoked responses of the young participants, in the ERPs and in the pupil dilation responses. Black line represents the across trials average response; green-purple lines represent single-trial responses.

**Supplementary Figure 2.**
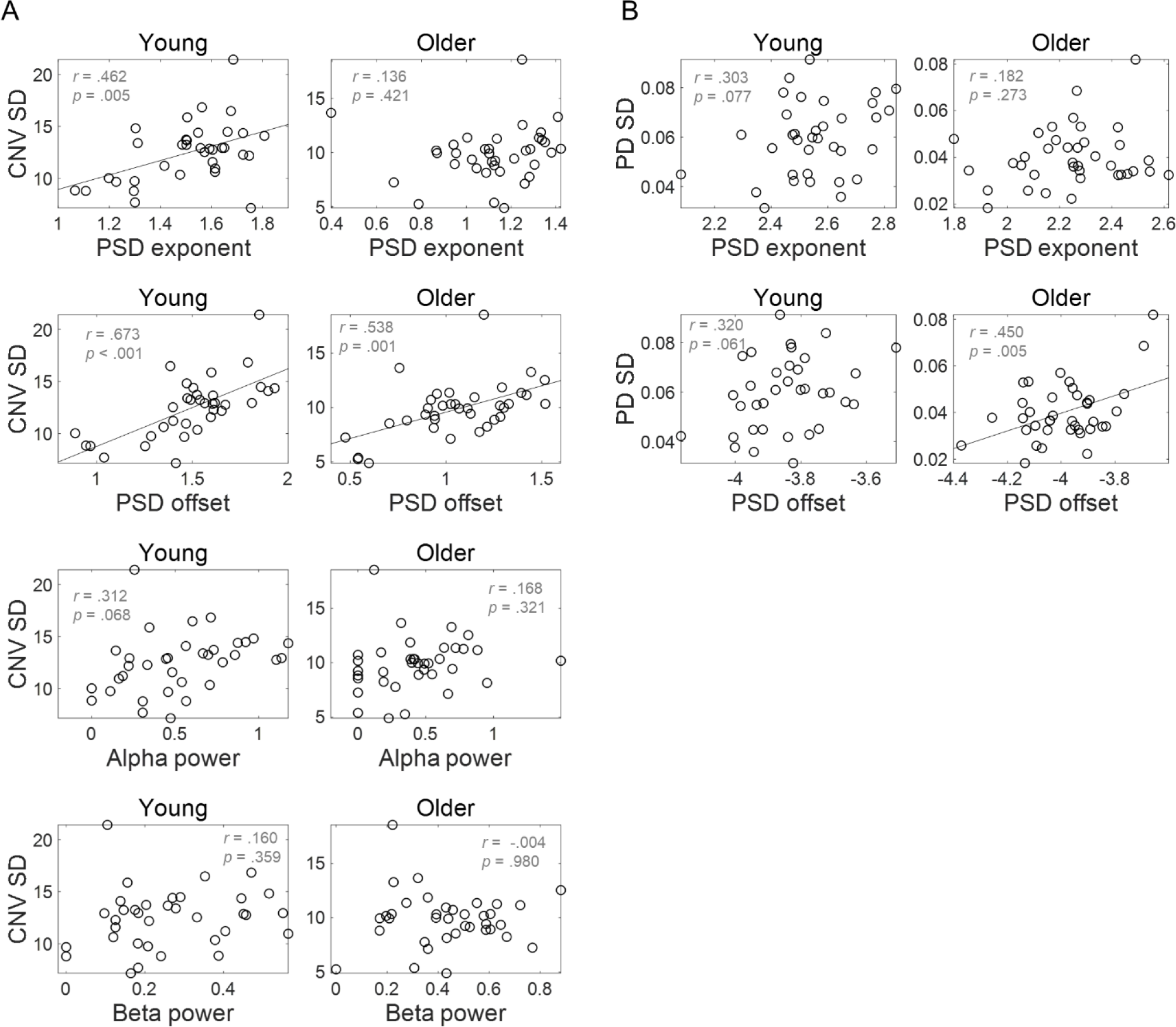
Variability in evoked responses correlates with the aperiodic parameters of the power spectral density (PSD) of the ongoing signals. (A) Across trials standard deviation (SD) of CNV amplitude plotted against (from top to bottom) exponent, offset, alpha power and beta power estimated from the PSDs of ongoing pre-stimulus EEG signal. CNV and ongoing signals measured at channel FCz. (B) Across trials SD of task-related pupil dilation (PD) response amplitude plotted against the aperiod parameters of the PSD of the ongoing pupil signal exponent and offset.

**Supplementary Figure 3.**
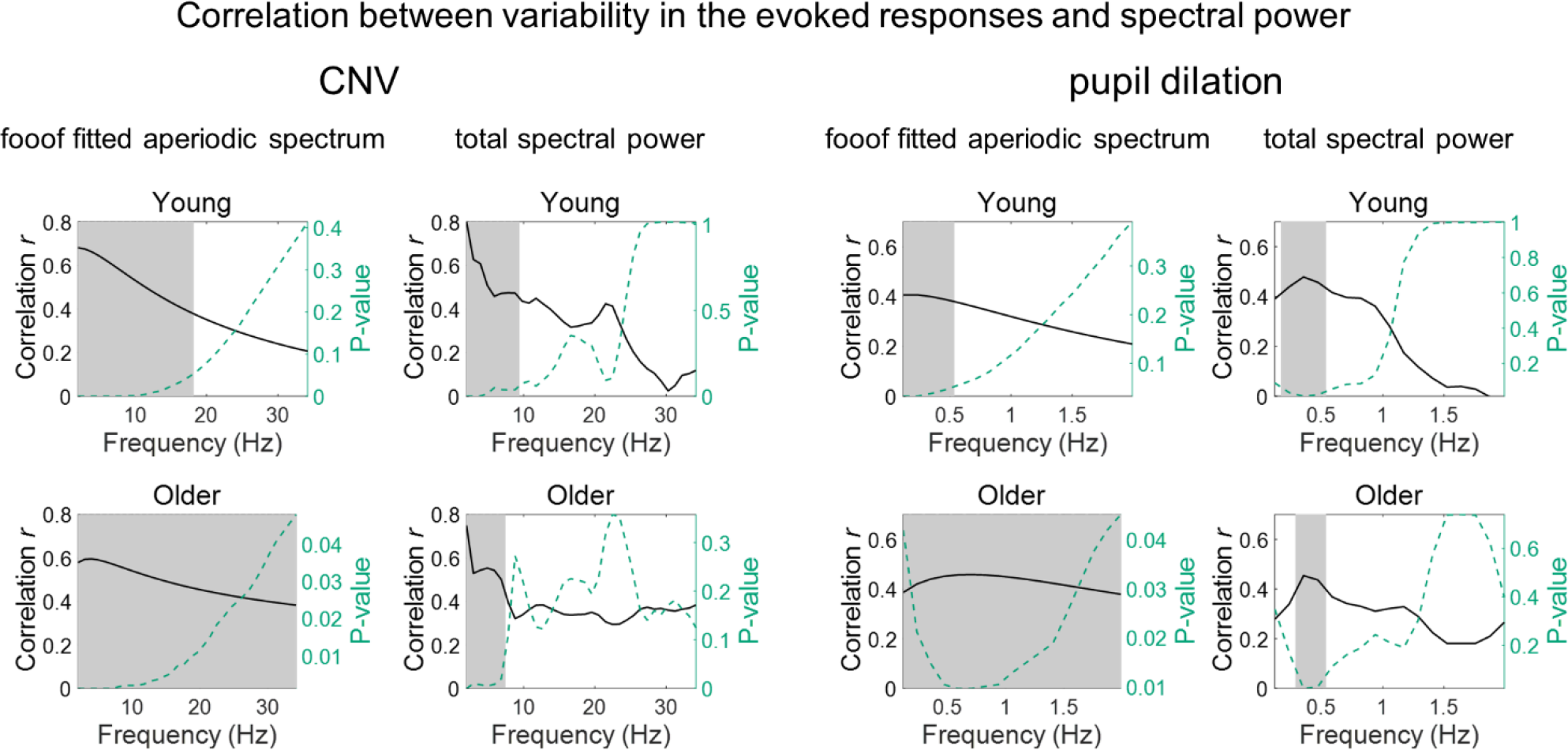
Pearson’s linear correlation coefficients between the power spectral density (PSD) at each frequency and the variability in the evoked responses of each participant. In the EEG signal, correlation between PSD and CNV variability was calculated in the FCz channel where the preparatory response presented highest amplitude. Grey background highlights the frequencies for which the correlation was significant. Permutation tests and the “max statistic” method were used for adjusting the *p*-values at each frequency for multiple comparisons.

**Supplementary Figure 4.**
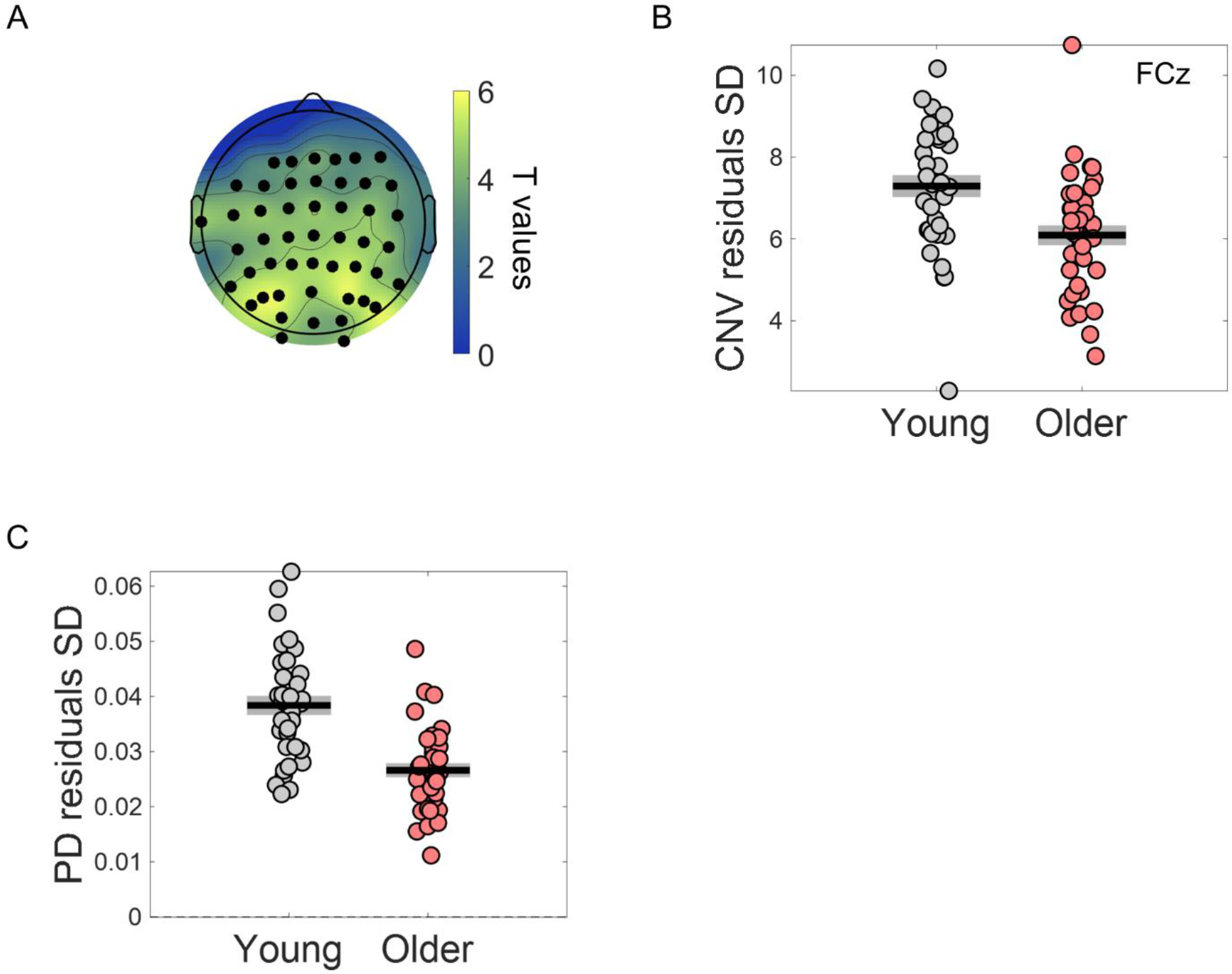
After regressing out the effect of ongoing slow fluctuations the variability of the evoked responses was still significantly reduced in the older group. (A) Scalp topographies of *t*-values from independent samples *t*-tests comparing the across trials standard deviation (SD) of CNV amplitude adjusted for the phase of ongoing EEG signal at cue-onset, in young and older individuals. Black circles highlight electrodes that show significant group differences. (B) CNV amplitude SD after regressing out the effect of the phase of ongoing EEG slow fluctuations at cue-onset measured at the EEG channel FCz. (C) Pupil dilation (PD) response SD after regressing out the effect of the phase of ongoing pupil slow fluctuations at cue-onset. Independent samples *t*-test revealed a significant effect of group *[t*_(71)_ = 5.70, *p* < .001]. (B and C) Circles = individual data points; black horizontal line = across participants mean; grey box = ± standard error of the mean across participants.

**Supplementary Figure 5.**
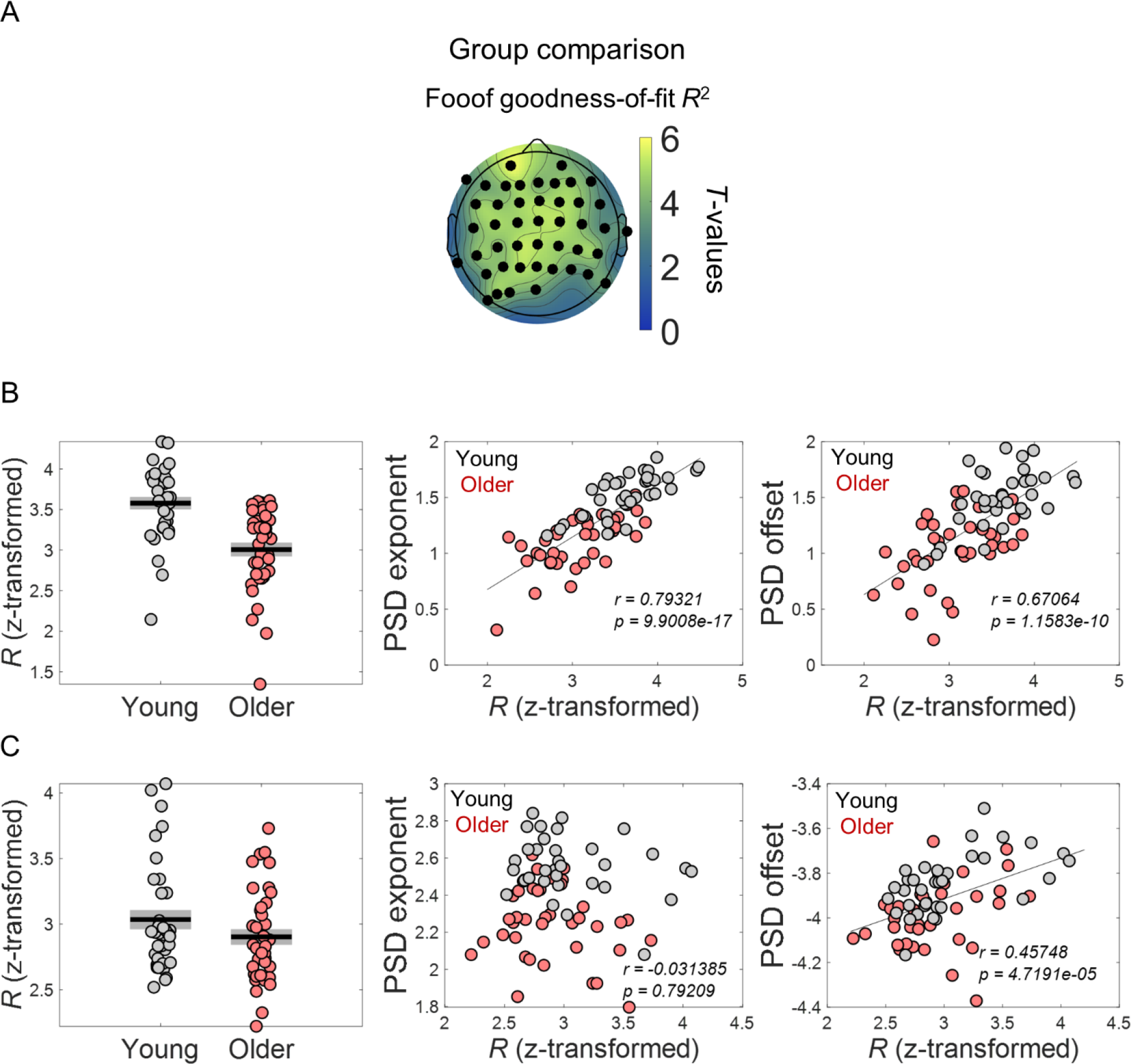
Goodness-of-fit of FOOOF model fitting to spectral data was significantly different across groups and correlated with the model aperiodic parameters, exponent and offset. (A) Scalp topographies of *t*-values from independent *t*-tests comparing goodness-of-fit *R* values (transformed into Fisher’s Z) across groups. Electrodes highlighted in black show statistically significant group differences after controlling for multiple comparisons. (B) Goodness-of-fit *R* (transformed into Fisher’s Z) of spectral data measured in electrode FCz (left). FCz exponent and offset parameters plotted against goodness-of-fit *R* (transformed into Fisher’s Z) (middle and right). (C) Goodness-of-fit *R* (transformed into Fisher’s Z) of spectral pupil data was also significantly reduced in the older group [independent *t*-test: *t*_(71)_ = 2.24, *p* = .028)]. In the pupil analyses, only the offset and not the exponent correlated with the *R* values (transformed into Fisher’s Z) (middle and right).

## Notes

https://openneuro.org/datasets/ds003690/versions/1.0.0

### Competing Interest Statement

The authors have declared no competing interest.

